# DAISM-DNN^XMBD^: Highly accurate cell type proportion estimation with *in silico* data augmentation and deep neural networks

**DOI:** 10.1101/2020.03.26.009308

**Authors:** Yating Lin, Haojun Li, Xu Xiao, Lei Zhang, Kejia Wang, Wenxian Yang, Rongshan Yu, Jiahuai Han

**Author notes:** These authors contribute equally (Yating Lin, Haojun Li).

## Abstract

Understanding the immune cell abundance of cancer and other disease-related tissues has an important role in guiding disease treatments. Computational cell type proportion estimation methods have been previously developed to derive such information from bulk RNA sequencing (RNA-seq) data. Unfortunately, our results show that the performance of these methods can be seriously plagued by the mismatch between training data and real-world data. To tackle this issue, we propose the DAISM-DNN^XMBD1^ pipeline that trains a deep neural network (DNN) with dataset-specific training data populated from a small number of calibrated samples using DAISM, a novel Data Augmentation method with an *In Silico* Mixing strategy. The evaluation results demonstrate that the DAISM-DNN pipeline outperforms other existing methods consistently and substantially for all the cell types under evaluation on real-world datasets.

## Introduction

It has been shown that the cellular composition of immune infiltrates in tumours is directly linked to tumour evolution and response to treatments^1, 2^. A high intratumoural infiltration of lymphocytes and dendritic cells is a favourable prognostic marker for cancer treatment^3, 4^, while a high stromal content of cancer-associated fibroblasts (CAFs) and M2 macrophages has been shown to be associated with poor outcomes^5, 6^. Particularly, recent progress in immunotherapy has led to durable clinical benefits, but only in a subpopulation of patients with “hot” tumour immune microenvironments that are characterized by a high infiltration of lymphocytes. Therefore, knowledge of the patient-specific immune cell proportion of solid tumours is invaluable in predicting disease progression or drug response as well as stratifying patients to select the most suitable treatment options.

In the past, fluorescence-activated cell sorting (FACS) and immunohistochemistry (IHC) were used as gold standards to measure the cellular components in a patient sample^7^. FACS requires a large number of cells, which limits its clinical applications. On the other hand, IHC only provides information on the cellular composition of a single biopsy slice which may not represent the full tumour microenvironment (TME) due to its heterogeneity. More recently, single cell RNA-seq (scRNA-seq) has emerged as a powerful technique to characterize cell types and states. However, the high costs of labour, reagents, and equipment of scRNA-seq at present restrain it from broad applications in routine clinical practice.

With the increasing availability of RNA quantification technologies, such as microarrays, high-throughput RNA-seq, and NanoString, the large-scale expression profiling of clinical samples has become feasible in routine clinical settings^8^. However, these methods only measure the average expression of genes from the heterogeneous samples in their entirety but do not provide detailed information on their cellular compositions. To bridge this gap, computational methods have been proposed to estimate individual cell type abundance from the bulk RNA data of heterogeneous tissues (Supplementary Table S1). In these methods, the abundance of each cell type from the mixed sample is quantified by aggregating the expression levels of the marker genes into an abundance score (MCP-counter^9^), by measuring the enrichment level of the marker genes using statistical analysis (xCell^10^), or by using computational deconvolution methods, such as least squares regression (quanTIseq^11^, EPIC^12^), support vector regression (SVR) (CIBERSORT^13^, CIBERSORTx^14^), or nonnegative matrix factorization (NMF)^15^, to derive an optimal dissection of the original sample based on a set of pre-identified cell type-specific expression signatures.

Undoubtedly, it is very challenging in practice for any of these computational methods to meet the rigid robustness and reliability requirements of biomedical or clinical studies over a broad range of sample types and conditions as well as sequencing technical platforms. For example, in deconvolution-based algorithms, it is expected that the cell type-specific expression signature should truly represent the expression characteristics of the underlying immune cells from the mixture samples. Unfortunately, the signature gene expression levels employed in existing methods are derived from either FACS-purified and *in vitro* differentiated or stimulated cell subsets or single-cell experiments. The application of antibodies, culture material or physical disassociation may affect the cell status, resulting in signatures that deviate from those of the actual cells *in vivo*. Moreover, technical and biological variations between RNA quantification experiments may introduce additional confounding factors that lead to sample or dataset-specific bias in cell type estimation. Similarly, marker gene expression aggregating methods such as MCP-counter require highly specific signatures with genes that are exclusively and stably expressed in certain cell types, which may not be possible for some immune cell lineages^16^.

Recently, the development of DNNs has granted computational power to resolve complex biological problems using data-driven approaches with the vast trove of data available from the biomedical research community powered by high-throughput genomic sequencing technologies^17, 18^. An application of DNN in cell type proportion estimation was proposed in Scaden^19^, where a neural network was trained on bulk RNA-seq data simulated from the scRNA-seq data of different immune cell types to predict cell type proportions from the bulk expression of cell mixtures. A DNN-based model could automatically create optimal features for cell fraction estimation during the training process, thus alleviating the need to generate reliable gene expression profile (GEP) matrices for different cell types. Moreover, it learns the potentially intricate nonlinear relationships between the gene expression composition and cell type proportions from training data, which are not possible to be captured by linear models used in other deconvolution algorithms. However, as the performance of DNN is still subject to the same statistical learning principle that test and train conditions must match, it is challenging for a DNN-based algorithm to deliver consistent performance under different experimental conditions unless sufficient ground truth data are available to train a specific predictive model for each distinct experimental condition. As a DNN model usually requires tens of thousands of training samples, the cost of implementing such a method would be prohibitive in practice.

To address these challenges, we developed the DAISM-DNN pipeline (Fig. 1) which consists of an *in silico* data augmentation method in collaboration with a DNN model to achieve robust and highly accurate cell type proportion estimation. The DAISM-DNN pipeline performs model training on a dataset augmented from a calibration dataset comprised of a small number of the actual data from the same batch of RNA-seq experiments, of which the ground truth cell type proportions are available for calibration. DAISM-DNN is able to deliver consistent cell type abundance profiling accuracies over different datasets. In addition, it is highly customizable and can be tailored to estimate the abundance of a large variety of cell types including those that are difficult for existing methods to estimate due to the lack of marker genes or GEP signature matrices and immune cells with overlapping markers or signature genes that are challenging, if not impossible, for existing methods^20, 21^.

**Figure 1:**
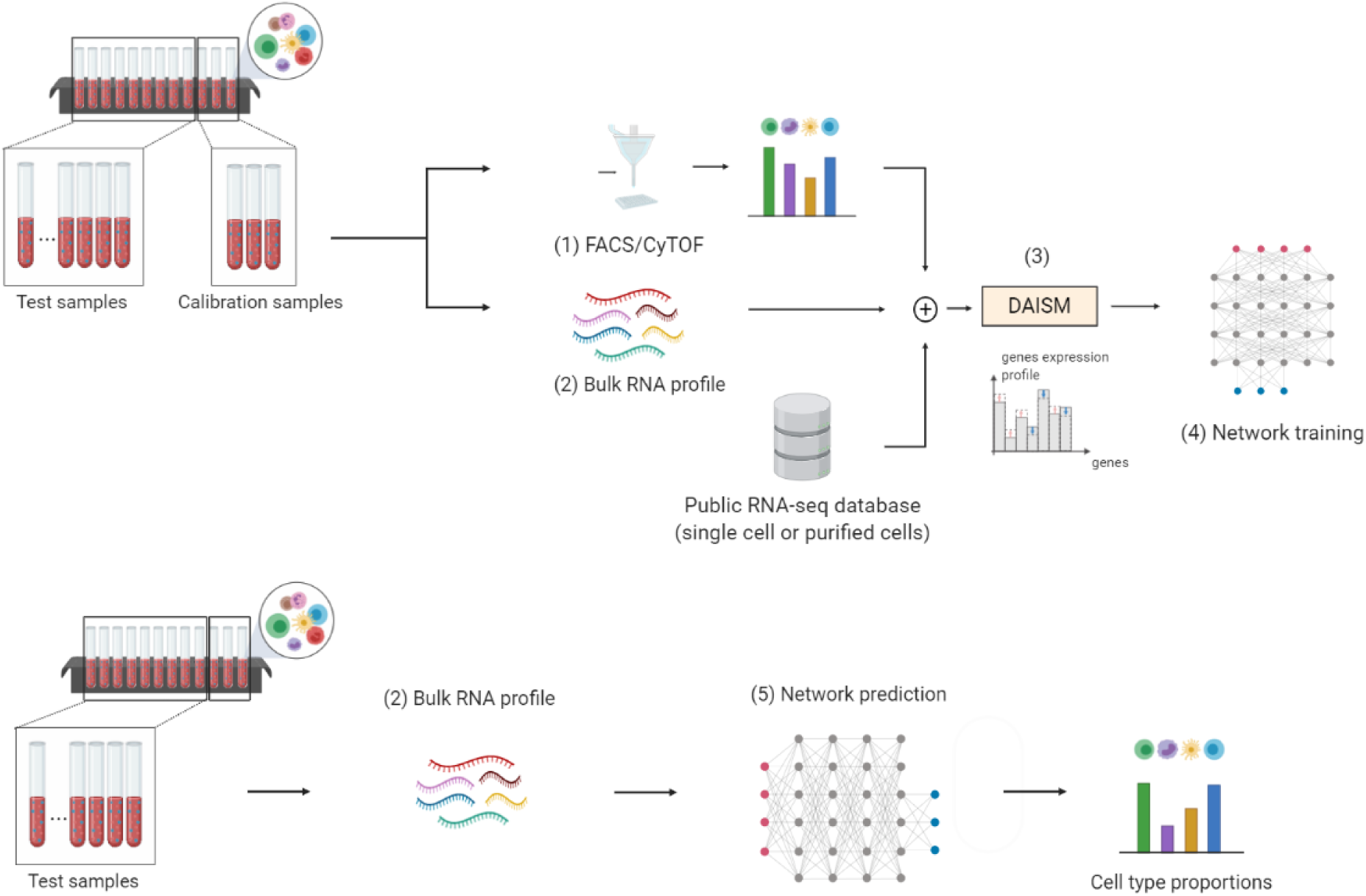
The DAISM-DNN pipeline. A typical DAISM-DNN workflow involves the following steps to perform cell type proportion estimation. (1) Measure the ground truth proportions of the cell types of interest in a small portion of calibration samples (e.g., 20∼40 samples) from the batch of samples to be evaluated. (2) Perform bulk RNA-seq on the calibration samples to obtain their expression profiles. (3) Perform data augmentation on the expression profiles of the calibration samples through *in silico* mixing with the RNA-seq data of purified cells or scRNA-seq data (DAISM). (4) Train a DNN using the augmented data. (5) Use the trained DNN model for cell type proportion estimation in the remaining samples with their bulk expression profiles. Steps (1)-(4) could be optional if the DNN model has already been trained for the given RNA-seq experimental conditions.

## Results

### There is no one-size-fits-all algorithm for cell type proportion estimation

We first evaluated nine state-of-the-art cell type proportion estimation algorithms, namely, CIBERSORT, CIBER-SORTx, EPIC, quanTIseq, MCP-counter, xCell, ABIS^20^, MuSiC^22^ and Scaden, on twelve independent real-world datasets (n = 685 total samples) acquired using different techniques or platforms (Supplementary Table S2) and three simulated datasets generated with scRNA-seq, purified bulk RNA-seq, and microarray respectively (Methods). Importantly, the methods under evaluation included the most recent developments to improve the cross-dataset robustness of cell type proportion estimation. For CIBERSORT, we further included four established basis signature matrices, namely, IRIS^23^, LM22^13^,TIL10^11^ and immunoStates^24^ in our evaluation. ImmunoStates^24^ used a basis matrix built using 6,160 samples with different disease states across 42 microarray platforms to mitigate the technical bias from different platforms. In MuSiC, the deconvolution algorithm further included appropriate weighting of genes showing cross-subject and cross-cell consistency to enable the transfer of cell type-specific expression information from one dataset to another. CIBERSORTx also implemented two batch correction modes (B-mode and S-mode) to reduce the potential bias from batch effects and we tested both modes.

Our results show that none of these methods were able to address the estimation bias problem to deliver consistently better results than others across multiple datasets. With regard to the overall prediction accuracy across all cell types and datasets, the DNN-based method (Scaden) achieved the highest average rank in terms of performance in RNA-seq data deconvolution while ABIS ranked first dealing with microarray data. However, the improvement over most other methods was not significant (Friedman test with post hoc two-tailed Nemenyi test, *α* = 0.05, Supplementary Fig. S1, Fig. S2a, b). In fact, as the performance of DNN is still subject to the same statistical learning principle that test and train conditions must match, Scaden, similar to other algorithms, showed inconsistent performance on different datasets. A t-distributed stochastic neighbour embedding (t-SNE) analysis of all test datasets demonstrated significant batch effects among those datasets and the difference among the testing samples was dominated by batch effects rather than cellular composition (Supplementary Fig. S2c). This result partially explains the inconsistent performance of the existing methods on different datasets and the challenge in developing a one-size-fits-most cell type proportion estimation method that performs uniformly well under different experimental conditions.

### DAISM-DNN enables accurate and robust cell proportion estimation

To overcome the aforementioned limitations, we developed DAISM, a data augmentation method, to produce dataset-specific training data for DNN model training, provided that a small number of RNA-seq data with ground truth cell type proportions from the same batch are available for calibration. DAISM generates a large amount of dataset-specific pseudo training data by performing *in silico* mixing of the calibration data with publicly available scRNA-seq data or RNA-seq data from purified cells at predefined ratios that are known to the training process. A DNN model that predicts the cell type proportions for the remaining samples is then trained using the DAISM-generated pseudo training data (Fig. 1).

We evaluated the performance of DNN models trained from DAISM-generated pseudo training data (DAISM-DNN) on the RNA-seq dataset SDY67. A total of 250 samples with ground truth proportions of five cell types (B cells, CD4 T cells, CD8 T cells, monocytes, and NK cells) from SDY67 were used for analysis in this paper. We used 200 randomly selected samples from SDY67 as calibration data, which were augmented with the scRNA-seq data of the five cell types to create the training data (Methods). The performance of DAISM-DNN was then tested on the remaining 50 samples that were not used in training. The experiment was repeated for 30 times independently. For comparison, we also employed other cell type proportion estimation algorithms on the same 50 testing samples. Overall, DAISM-DNN outperformed all other algorithms by a significant margin from 30 repeated experiments for all the cell types under evaluation (Fig. 2 and Supplementary Fig. S3). When evaluated by the average per-cell-type Pearson correlation between the predicted and ground truth cell proportions, DAISM-DNN achieved the highest correlation, followed by Scaden (boxplot in Fig. 2b; results after 30 bootstrapping experiments). In addition, DAISM-DNN had the lowest root mean square error (RMSE) and the highest Lin’s concordance correlation coefficient (CCC), followed by ABIS (Fig. 2b). We replaced the scRNA-seq data with the RNA-seq data of purified cells to generate training data with DAISM and did not find a significant difference in performance between the DNN models derived from these two approaches (DAISM-RNA vs DAISM-scRNA; Supplementary Fig. S4).

**Figure 2:**
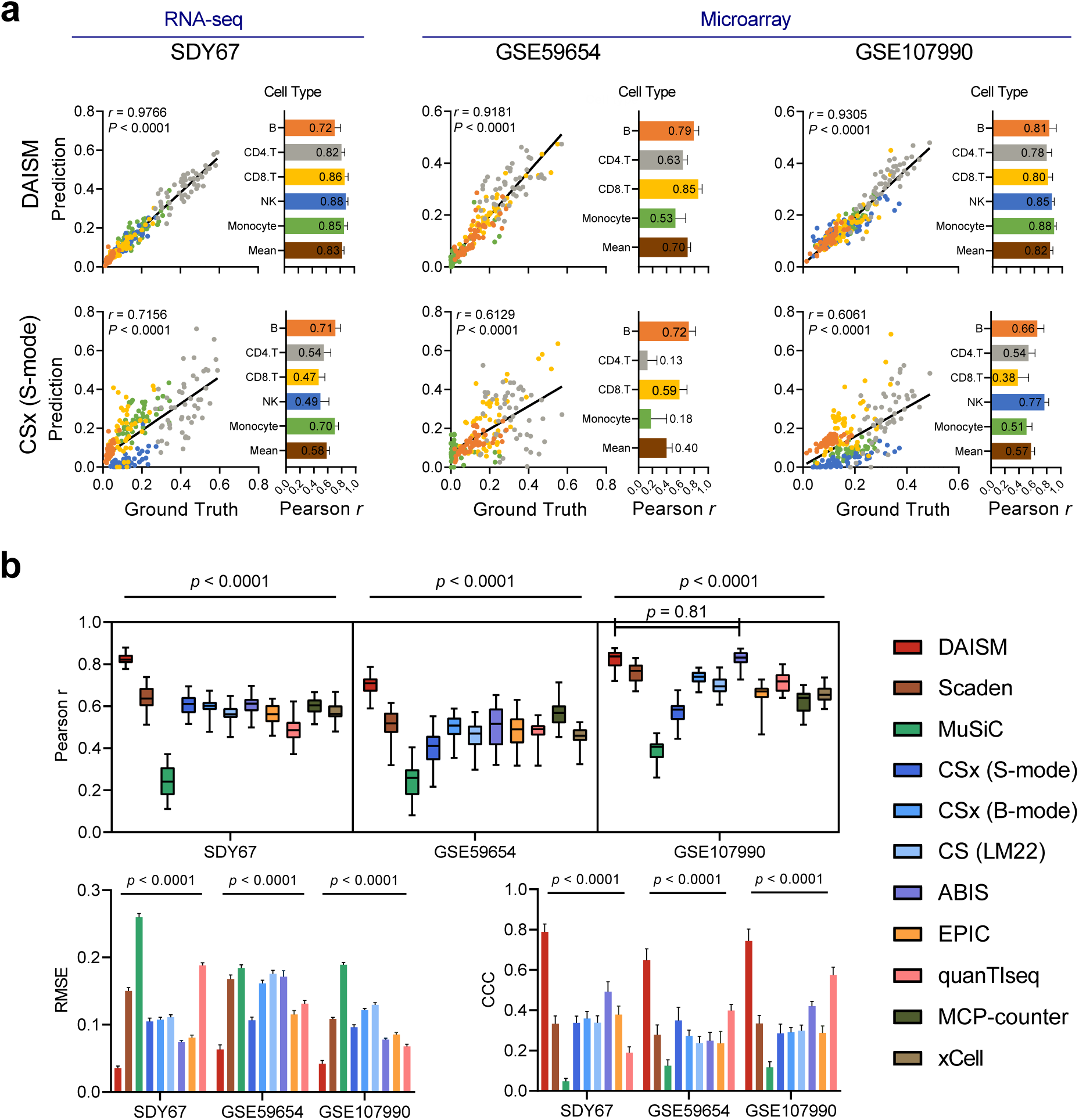
Performance of different algorithms on datasets SDY67, GSE59654, and GSE107990. (a) Scatter plots of ground truth fractions (x-axis) and predicted cell fractions (y-axis) for DAISM-DNN and CIBERSORTx (S-mode). The bar plot shows the Pearson correlation for each cell type in 30 bootstrapping experiments. The value in bar plots indicates the mean value of 30 experiments. (b) Boxplot of the mean of per-cell-type Pearson correlations for 11 methods, and bar plots of RMSE (right) and CCC (left) for 9 methods. All data in bar plots are presented as the mean±SD. Note that RMSE and CCC are not suitable for evaluating the two marker-based methods, MCP-counter and xCell. Two-sided paired Student’s t tests were used for comparing DAISM-DNN with other methods.

We further tested DAISM-DNN on two microarray datasets GSE59654 and GSE107990 with 153 and 164 samples respectively. Similarly, we randomly selected 50 samples as test data, and the remaining samples were augmented with the scRNA-seq data of the respective cell types to generate the training dataset for DAISM-DNN (Methods). The bootstrapping results showed that DAISM-DNN outperformed the other algorithms by a significant margin for all the cell types under evaluation (Fig. 2 and Supplementary Fig. S3), except for GSE107990, where the difference between ABIS and DAISM-DNN was not significant.

We extended our evaluation to a fine-grain cell population of 14 cell types: naive B cells, memory B cells, naive CD4 T cells, memory CD4 T cells, regulatory T cells, naive CD8 T cells, memory CD8 T cells, monocytes, NK cells, macrophages, neutrophils, myeloid dendritic cells (mDCs), fibroblasts, and endothelial cells. The comparison only performed on nine cell types on these three validation data, because of the lack of ground truth data for some fine-grain cell types (i.e., regulatory T cells, macrophages, neutrophils, fibroblasts, and endothelial cells). The results were compared with those of CIBERSORTx (S-mode and B-mode), ABIS, and xCell, which are also able to produce estimations of fine-grain cell type proportions. Comparisons were only performed on cell types where ground truth cell type proportion information was available for each dataset (Supplementary Fig. S5). The results indicate clear advantages of DAISM-DNN over traditional methods not only in overall performance but also for all individual cell types and datasets in terms of the Pearson correlation (Supplementary Fig. S5b), RMSE, and CCC (Supplementary Fig. S5c).

To understand if the performance gain of DAISM-DNN indeed comes from the data-specific training set generated using DAISM, we generated *in silico* mixed training data using DAISM with calibration data from SDY67 as well as the direct mixing of RNA-seq data from sorted cells or scRNA-seq data of selected cell types (Methods). All the *in silico* mixed data from different mixing strategies followed the same cell type proportions. The t-SNE plot revealed highly distinct clusters of these datasets. Importantly, only the clusters of the DAISM-mixed dataset strongly overlapped with SDY67 while the clusters from the remaining datasets showed a clear gap from SDY67, demonstrating strong batch effects between them and the real samples (Fig. 3). We also used a combination of simulated mixtures and real data with cell fraction information as suggested in Menden et al.^19^ to train DNN models. We integrated five RNA-seq real-world data, including 200 samples excluded for testing of SDY67, with simulated data generated by scRNA-seq data respectively. The training data size kept the same in these separate trainings. Obviously performance increases were observed only when seamlessly combining calibration samples with simulated training dataset. Furthermore, DNN models trained with DAISM-mixed training datasets achieved significantly better performance than those trained with other *in silico* training data (Fig. 3b), demonstrating the critical role of training data in determining the performance of DNN-based models and the effectiveness of DAISM in creating a training dataset that matches the intrinsic distributions of the real-life data to enable highly accurate cell type proportion estimation.

**Figure 3:**
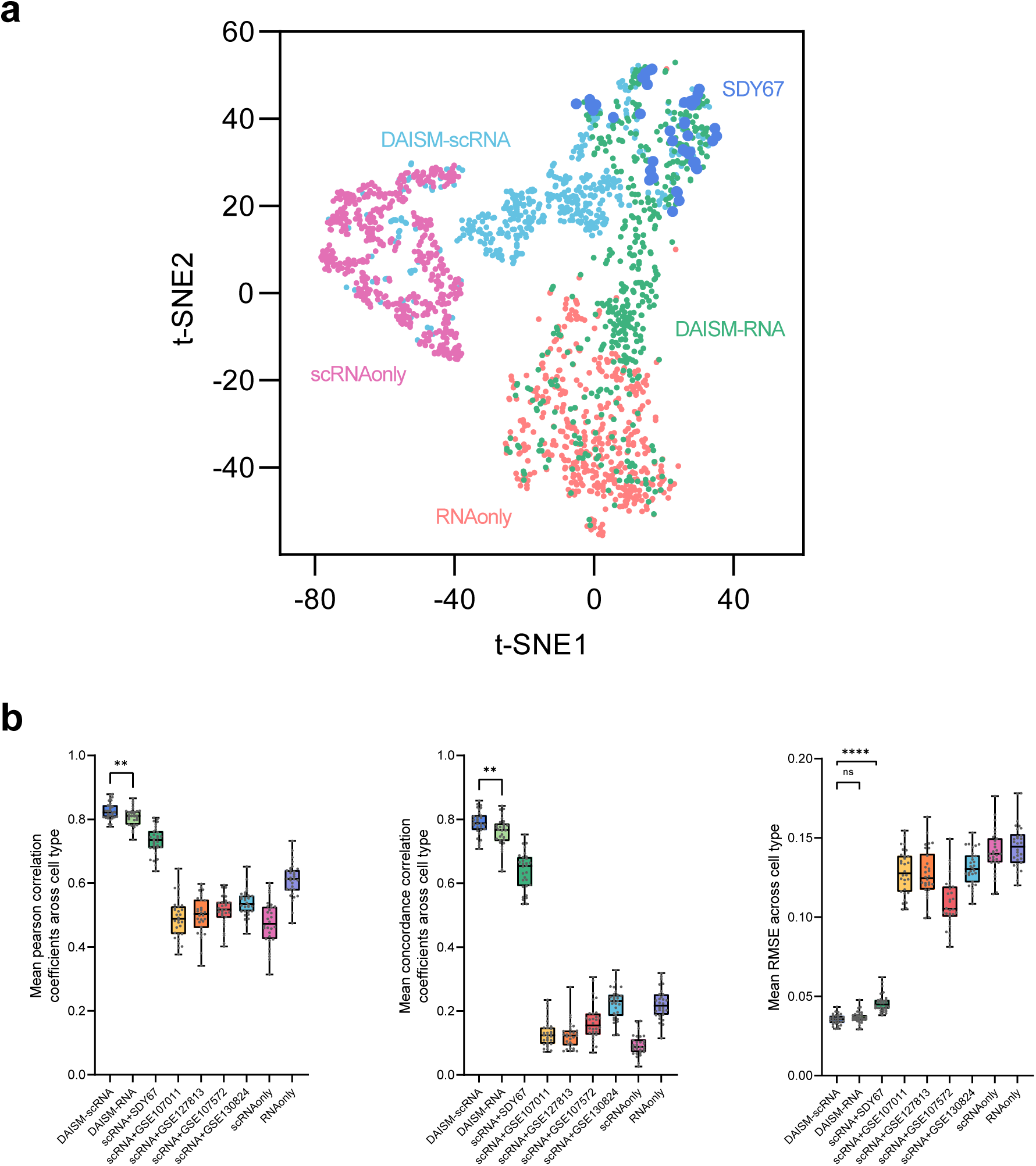
Performance of DNN on different training datasets. (a) The t-SNE projection of the SDY67 dataset (*n* = 50 samples) and training datasets (*n* = 500 samples per platform) constructed by different strategies and purified samples from different platforms, coloured according to training datasets. (b) The cell-type proportion estimation performance on SDY67 was evaluated using Pearson correlation (left), CCC (middle) and RMSE (right). Models trained by four training datasets. \*\*\*\**P <* 0.0001,\*\**P <* 0.01, two-sided paired Student’s t-test.

### Applicability of DAISM-DNN in real-life biomedical experiments

A limitation of DAISM-DNN is that it requires calibration samples to train the dataset-specific models, which may not always be available. On the other hand, it is common that in many scenarios, rigorous standard operation procedure (SOP) and quality control are enforced to derive consistent gene expression quantification results across experiments, such as defined by the gene expression based biomarker Oncotype DX ^25, 26^. In cases where strict SOPs are enforced, it is possible to pre-train a generic DAISM-DNN model that can be used across different batches without the need for retraining every time. To verify this concept, we generated a validation dataset comprising 36 human peripheral blood mononuclear cells (PBMCs) samples assayed by the same RNA-seq panel in two separate batches (Methods). The first batch consists of 30 samples which were used as calibration samples to generate DAISM-mixed training dataset, while the other batch consists of 6 samples for testing. The ground truth cell-type proportions of both batches were established using mass cytometry (CyTOF, see Methods).

To generate the DAISM-mixed data for training in this study, we used CITE-seq^27^ data, which provide single-cell transcriptome and surface proteins simultaneously, from two public CITE-seq datasets (PBMC5k and PBMC10k) for augmentation. Clusters of different cell types were identified seperately for both CyTOF and CITE-seq datasets through meta-clustering on 11 surface marker proteins in common of these two datasets, and manually annotated based on canonical marker expression patterns consistent with known immune cell types. These clusters were further pairwise linked according to Pearson correlation of normalized mean marker expression of each cluster to identify matching populations between them (Fig S7, Fig S8b; Methods). It can be seen from the results that with strict SOP, it is possible to minimize the technical variance of gene expression results between batches (Supplementary Fig S8a), and enable more stable, robust and accurate cell type proportion estimation compared to other established methods through a pre-established DAISM-DNN model trained on different batches (Fig S7c).

Finally, to identify the impact of the number of calibration samples on the performance of DAISM, we tested the cell proportion estimation performance of DAISM-DNN with respect to the number of calibration samples used in creating the augmented training data. We found that in general, the estimation accuracy improved with an increasing number of calibration samples used in creating the *in silico* mixed training data (Supplementary Fig. S6). When evaluated by CCC or RMSE, which requires that the predicted cell fractions follow the real fractions in terms of absolute numbers, the estimation performance improved dramatically when the number of real samples used in *in silico* mixing increased from zero to 20∼40. Beyond that, the rate of improvement slowed down significantly with more calibration samples, suggesting that a calibration dataset of reasonable sizes would be sufficient for DAISM to generate dataset-specific training data to train a DNN model with satisfactory cell type proportion estimation performance.

### DNN reveals biologically meaningful markers that cannot be leveraged by traditional deconvolution methods

We used the SHAP values^28^, which explain the output of a predictive model as the sum of the effects from each individual input feature introduced into a conditional expectation, to rank the importance of individual genes for the prediction of different cell types. We calculated the average absolute SHAP values of all genes for all samples of SDY67 using DeepSHAP^28^ for the proportion prediction of B cells, NK cells, monocytes, CD8 T cells and CD4 T cells. A clear long tail of the distribution of the SHAP values can be observed, suggesting that a small number of the “marker genes” play an essential role in the cell proportion prediction results (Fig. 4a). A correlation study of the top 50 genes ranked by SHAP values for each cell type showed that most of these genes were positively correlated with the abundances of the corresponding cell types, and some of these positively correlated essential genes identified by SHAP value analysis are indeed marker genes of the corresponding cell types reported in the literature (Fig. 4b).

**Figure 4:**
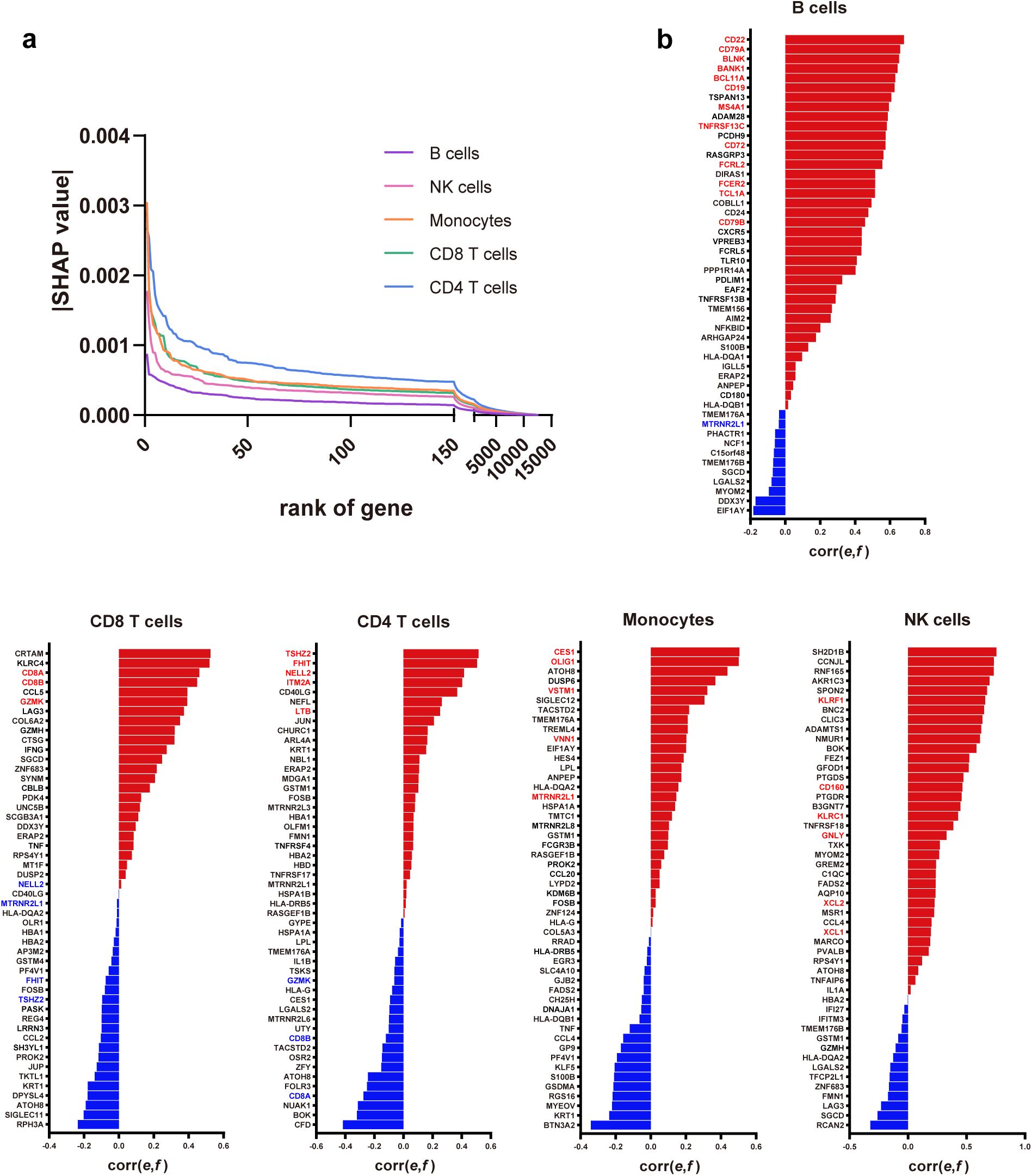
Interpretation of DAISM-DNN using SHAP on the SDY67 dataset. (a) Distribution of absolute SHAP values for the rank of important feature genes across coarse-grain cell types. (b) Pearson correlation between the expression level *e* and cell type fraction *f* of the top 50 important features selected by SHAP for each cell type. Red bars represent positive correlations and blue bars represent negative correlations. For each cell type, the biological markers of corresponding cell types are labeled in red, and markers of other cell types are labeled in blue.

Interestingly, the SHAP-identified important genes also included those that are highly expressed in more than one cell lineage. For example, SH2D1B and CCNJL, the top-ranked genes selected by DNN for NK cells, are also highly expressed in nonclassical monocytes and neutrophils. These genes would not usually be selected during the gene selection process in traditional marker or signature-based methods due to the lack of specificity. Moreover, some genes deemed essential features by SHAP value analysis were only weakly or even negatively correlated with the target cell type of the trained prediction models. Closer examination showed that some of those negatively correlated genes are in fact marker genes of the closely related cell types from the same lineage as the target cell types (e.g., the marker genes CD8A, CD8B, and GZMK for CD8 T cells were identified as essential genes for the prediction of CD4 T cell proportions), suggesting that the DNNs learned to use these genes as negative controls to better differentiate highly confused cells to achieve better prediction through the training process.

## Discussion

Understanding the cellular heterogeneity in disease-related tissues is essential for the identification of cellular targets for treatments. To this end, computational methods have been developed to quantify cell type compositions from the GEPs of bulk samples, thus allowing the elucidation of cell type contributions to disease from highly available disease-related bulk RNA-seq data. Existing deconvolution methods rely on pre-selected cell type-specific marker genes or signatures based on cell type-specific gene expression, which could be derived from existing RNA-seq datasets of single cells or purified cell lines of target cells. The accuracy of these methods is therefore subject to the effectiveness of the selected GEPs to represent different bulk RNA-seq datasets under testing. Unfortunately, due to the presence of strong non-biological cross-platform variations, the performance of such methods could fluctuate greatly when applied to different datasets even with the latest advancements in batch normalization^14^ or platform-agnostic signature designs^22, 24^.

We developed DAISM-DNN to meet the challenge of accurate cell type proportion quantification for bulk tissues using GEPs derived from disparate sources. One of the key features that differentiates our method from previous works is that we used a DNN-based, data-driven approach that is free from manually curated marker genes or expression signatures. By learning directly from the data, DAISM-DNN not only discovers new features that were previously unrevealed from conventional methods, but also leverages their intricate interactions with the target phenotypes to achieve accurate prediction results, which is impossible when shallow models are used.

Another key feature of DAISM-DNN is that instead of relying on normalization or platform-agnostic reference profiles to overcome the cross-platform variation problem, DAISM-DNN builds a dataset-specific prediction model from a small number of calibration samples from the testing dataset, thus fundamentally avoiding the problem. This requirement may seem stringent and restrictive as the calibration samples need to be sorted to establish the ground truth cell type proportions. However, our results indicated that only a small number of calibration samples are needed to train the prediction model thanks to the DAISM data augmentation strategy. More importantly, we have also shown that with stringent SOPs in the overall experimental procedure, such as those being practiced for GEP based assays for clinical usage^26^, it is possible to create a “train once, reuse many times” assay-specific DAISM-DNN model for data generated under the same or similar experimental conditions. Overall, DAISM-DNN is particularly suitable for large cohort studies or routine clinical applications of which highly reliable and accurate cell fraction information is expected, and relatively stable RNA-seq experimental conditions are involved.

Deep learning models are often deemed black boxes. Although they provide accurate prediction results, the learned relationships between features and results are typically hard to interpret. By assessing our trained DNN models with regard to the determinants of their predictions with SHAP values, we found that different genes differ remarkably in terms of their contributions to the cell type proportion estimation. Importantly, we found that each individual cell type has a distinct group of genes that are deemed essential to the prediction results based on their SHAP values, and many of these essential genes coincide with the marker genes of the target cell type identified in the literature. Moreover, SHAP analysis also revealed that many essential features selected by the DNN models are in fact marker genes of closely related cell types (e.g., cell types from the same lineage of the target cell type), which are apparently used by the DNN models as negative controls to differentiate highly confused cell types. These results highlight that our model indeed learned the intricate relationships between gene expression and the cellular composition of the bulk samples that are difficult to leverage in shallow models.

Finally, despite the success of deep learning, people find it challenging to apply such models in genomic studies in a supervised learning setup due to the scarcity of training samples. The data augmentation strategy as in DAISM provides a broadly applicable framework to create a large amount of *in silico* mixed artificial training data from a small amount of real-life samples with the aid of the increasing availability of scRNA-seq datasets or other datasets that provide comprehensive characteristic maps of different cell types from single-cell analyses. We hypothesize that the algorithmic principles underlying DAISM could be generalized to other gene expression analysis tasks that are currently incompatible with deep learning due to training data limitations. Future studies are warranted to demonstrate these possibilities.

## Methods

### Data augmentation through *in silico* mixing (DAISM)

Deep learning-based approaches require a large amount of training data. In general, existing data from real tissue samples with known fractions of cell types and gene expression levels could be insufficient to use as a training set. In this regard, we extracted a small number of real-life samples with ground truth cell type proportions to use as a calibration dataset, and applied the DAISM strategy to create a large number of *in silico* mixing samples from this calibration dataset.

The expression profile of a DAISM-generated (i.e., *in silico* mixed) sample is calculated as follows. First, we generate a random variable *r* with uniform distribution between 0 and 1 to determine the fraction of the real sample in the mixed sample, and *C* random variables with Dirichlet distribution *p*_*k*_, (*k* = 1, …, *C*) such that 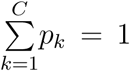 to determine the fractions of the immune cells in the mixed sample, where *C* is the number of cell types. The expression profile of the final mixed sample ***e*** is then calculated as:

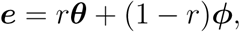

where ***θ*** is the expression profile of a real-life sample randomly selected from the calibration dataset as a seed sample for this *in silico* mixed sample, and ***ϕ*** is the aggregated expression of single cell samples or purified samples used for data augmentation. When using scRNA-seq dataset for data augmentation (DAISM-scRNA), we have

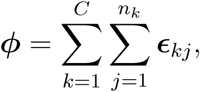

where *n*_*k*_ = 500 · *p*_*k*_ is the number of cells of type *k* extracted randomly from scRNA-seq datasets for mixing, and ***ϵ***_*kj*_ denote their expression profiles. Note that ***ϕ*** is further TPM-normalized before mixing. When using RNA-seq or microarray data from purified cells (DAISM-RNA), we have

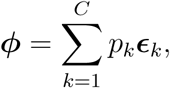

where ***ϵ***_*k*_ is the expression profile of a randomly selected purified sample of cell type *k* from the respective RNA-seq dataset. Once the expression profile of the *in silico* sample is created, its “ground truth” cell fractions can be calculated as:

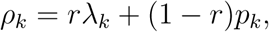

where *ρ*_*k*_ is the fraction of cell type *k* in the *in silico* mixed sample, *λ*_*k*_ is the ground truth fraction of cell type *k* in the selected seed sample which is known *a priori* through experiments, e.g., flow cytometry analysis.

### The DAISM-DNN pipeline

We trained deep feed-forward, fully-connected neural networks (multilayer perceptron networks) on the *in silico* mixed training data obtained by DAISM to predict the cell fractions from bulk expression data. The network consists of one input layer, five fully-connected hidden layers and one output layer, implemented with PyTorch (v1.0.1) in Python (v3.7.3). As a DNN can fit a large feature space with a large number of parameters (i.e., connection weights), we did not perform feature selection in advance. Instead, we used all the genes that were present in both the training and testing datasets as input to the neural network. Moreover, the expression profile of each sample was log2-transformed, and scaled to the range of [0,1] through min-max scaling before input to DNN:

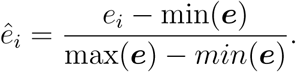

Here, *e*_*i*_ is the log2-transformed expression level of gene *i*, ***e*** is the vector of the log2-transformed expression levels of all genes of a sample, and *ê*_*i*_ is the vector of min-max scaled values.

The network was trained using the back-propagation algorithm with randomly initialized network parameters. The mean square error (MSE) between the actual and predicted absolute cell fractions was used as the loss function. The optimization algorithm Adam was used with an initial learning rate of 1e-4. During the training process, the training set was randomly divided into minibatches with a batch size of 64. When the average MSE of all mini-batches in the current epoch was higher than that of the last epoch, the learning rate was multiplied by an attenuation coefficient until a minimum of 1e-5 was reached to avoid training noise from excessively large learning rates when the network converged to steady state. We randomly split the training set and the validation set at a ratio of 80:20. Early-stopping strategy was adopted to stop training when the validation error did not decrease in 10 epochs, and the model producing the best results on the validation set during training was selected as the final model for prediction.

### RNA-seq datasets of purified cells

For the RNA-seq data of purified immune cells to serve as augmentation data, we used 1533 purified cell samples of the eight immune cell types (B cells, CD4 T cells, CD8 T cells, monocytes, NK cells, neutrophils, endothelium, and fibroblast) in this study (Supplementary Table S3). The raw FASTQ reads were downloaded from the NCBI website. Transcription and gene-level expression quantification were performed using Salmon^29^ (version 0.11.3) with Gencode v29 after quality control of FASTQ reads using fastp^30^. All the software tools were used with their default parameters. Transcripts per million (TPM) normalization was then performed on all the samples.

### scRNA-seq datasets

For the scRNA-seq data of different immune cell types, we downloaded a scRNA-seq dataset of PBMCs from patient blood samples from the 10X Genomics website (https://support.10xgenomics.com/single-cell-gene-expression/datasets, “8k PBMCs from a Healthy Donor”), denoted as PBMC8k. PBMC8k was sequenced on Illumina Hiseq4000 with approximately 92,000 reads per cell, and 8,381 cells were detected in total. We used this dataset for augmentation in both coarse-grain and fine-grain deconvolution. The raw scRNA-seq reads were aligned to the GRCh38 reference genome and quantified by Cell Ranger^31^ (10X Genomics version 2.1.0). The resulting expression matrix was then processed using Seurat^32^ (v. 3.1.1). First, cells with less than 500 genes or greater than 10% mitochondrial RNA content and genes expressed in less than 5 cells were excluded from analysis. Then, cells with abnormally high gene counts were considered as cell doublets and were excluded from further analysis. The raw unique molecular identifier (UMI) counts were log-normalized and the top 2000 highly variable genes were called based on the average expression (between 0.0125 and 3) and average dispersion (*>* 0.5). Principal component analysis (PCA) was performed on the highly variable genes to further reduce the dimensionality of the data. Finally, clusters were identified using the shared nearest neighbor (SNN)-based clustering algorithm on the basis of the first 20 principal components with an appropriate resolution.

The identified clusters were annotated on the basis of marker genes’ expression levels. The marker genes were obtained from the CellMarker database^33^ for the target cell types in peripheral blood, specifically, CD4 for CD4 T cells, CD8A and CD8B for CD8 T cells, MS4A1 and CD79A for B cells, CD14 and FCGR3A for monocytes, GNLY for NK cells, FLT3 and FCER1A for dendritic cells. Cell types were identified manually by checking if the respective marker genes were highly differentially expressed in each cluster. The clusters without high expression on the selected marker genes or with high expression on the marker genes of other cell types were grouped into the “unknown” type.

For fine-grain deconvolution, PBMC8k was further clsutered into finer groups based on the major cell types and served as augmentation data. B cells were subclusterd into naive B cells and memory B cells. CD4 T cells and CD8 T cells were further grouped into naive CD4 T cells, memory CD4 T cells, naive CD8 T cells and memory CD8 T cells, respectively. Myeloid dendritic cells and monocytes came from monocyte lineages.

### CITE-seq datasets

In deconvolving the in-house validation dataset, we used two CITE-seq PBMC datasets which provide read counts of both mRNAs and cell surface proteins as augmentation data. The datasets were downloaded from 10X Genomics website (PBMC5k from https://support._10xgenomics.com/single-cell-gene-expression/datasets/3.0.2/5k_pbmc_protein_v3, PBMC10k from https://support.10xgenomics.com/single-cell-gene-expression/datasets/3.0.0/pbmc_10k_protein_v3). The PBMCs from a healthy donor were stained with TotalSeq-B antibodies (BioLegend) in both datasets for identification of cell surface proteins, and were sequenced by Illumina NovaSeq for scRNA-seq. For mRNA profiles in CITE-seq, the same preprocessing steps for scRNA-seq data were applied for quality control. The protein counts were pre-processed with the centered-log-ratio (CLR) normalization^27^ prior to clustering.

### Datasets for benchmarking

We used eleven public microarray and bulk RNA-seq datasets to evaluate the performance of different cell type proportion estimation methods, of which the expression data and the corresponding ground truth cell fractions were publicly available with reference to the original publications and used accordingly in our benchmarking tests. The cell fractions for SDY67, GSE107990 and GSE59654 were taken from the supplementary materials of the publication^20^. A table listing details on all datasets including references to the original publications can be found in Supplementary Table S2.

We also generated three datasets of simulated mixtures using single cell samples from PBMC8k, purified bulk RNA-seq samples and microarray samples respectively. Each dataset contains 50 samples.

### In-house PBMC validation samples

To further evaluate the validity of our method, we generated an in-house validation dataset of human PBMCs from a cohort of drug addicts and healthy donors. A total of 36 PBMC samples were isolated from whole blood by Ficoll density gradient centrifugation and stored in liquid nitrogen with frozen solution (90% FBS and 10% DMSO). Cryopreserved PBMCs were thawed with pre-warm complete RPMI 1640 medium (RPMI1640, 10% FBS) containing 25U/ml benzonase. Cells from each sample were washed, counted, adjusted to 2 × 10^6^ living cells/stain and transfered to a new 5ml polystyrene round-bottom tube. Simultaneously, 1 × 10^5^ living cells/sample were washed with PBS and quickly frozen in liquid nitrogen for RNA-Seq.

### Mass cytometry of validation samples

Isotope-labeled antibodies were purchased from PLT Tech Inc. (Hangzhou, China), where antibodies conjugation and testing were performed. After thawing and preprocessing, PBMC samples were stained by surface antibodies (Supplementary Table S4) to 2 × 10^6^ cells for 30 minutes at RT. All samples were then washed with Maxpar Cell Staining Buffer and incubated in Nuclear Antigen Staining Buffer for 30 minutes at RT. Then, samples were washed and stained by intracellular antibody cocktail for 30-45 minutes at RT. Subsequently, cells were washed and incubated in Ir intercalation solution overnight at 4°C. Immediately prior to data acquisition, cells were washed, and Maxpar water with EQ Beads were added to adjust cell concentration to 2.5 ∼ 5 × 10^4^/ml. All samples were acquired on a CyTOF2 mass cytometer (Fluidigm, Helios) at an event rate of 200-500 cells per second.

### RNA-seq of validation samples

RNA was extracted using RNeasy Mini Kit (QIAGEN) following the manufacturer’s protocol. Barcoded RNA libraries were generated and captured by a customized Master panel. All libraries were sequenced on the Illumina NovaSeq 6000 platform with 2×150 bp paired-end reads. For processing RNA-seq data, quality assessment was carried out using in-house script FormatFastQ (v2.4.0). Alignment to targeted genes from GenCode hg37 was performed using STAR (v2.7.2b) and gene counts were quantified using RSEM (v1.3.3).

### CyTOF data analysis

CyTOF data were normalized by EQ-bead normalization in the CyTOF2 equipment, and uploaded to Cytobank (https://community.cytobank.org/) for data cleaning, doublets and dead cells removal. We removed EQ beads, used channel DNA2 and Event length to exclude aggregated cells, and used channels DNA2 and Rh103 to select alive cells. The results are then exported as .fcs files for analysis. Data were scaled with arcsinh-transformation and further analyzed in R (v3.6.3). Single alive cells were first clustered using FlowSOM^34^ (R package, v.1.18.0) which used self-organizing maps for high-dimensional data reduction. Subsequently, we used Phenograph (R package, v0.99.1), a graph-based community detection method using the Louvain algorithm, for second clustering on the groups from FlowSOM. After every single cell was assigned to a cluster, we manually annotated each cluster based on its marker expression pattern compared with patterns of known immune cell types.

### Mapping between CyTOF and CITE-seq cell clusters

To find proper augmentation data for deconvolving the in-house validation dataset, we first performed clustering on normalized protein profiles of CyTOF and CITE-seq respectively using both FlowSOM and Phenograph. Eleven surface markers in common were used for clustering: CD3, CD4, CD8a, CD14, CD16, CD56, CD19, CD25, CD45RA, CD45RO, CD127. Then the Pearson correlation between each cluster of CyTOF and CITE-seq data was calculated based on the mean values of marker expressions. For each CyTOF cluster, we identified the best-matching cluster of CITE-seq according to the correlation between two clusters. We allowed one-to-many mapping in pairwise linking. After building these pairwise constraints, we further manually annotated these clusters based on similar typical marker expression patterns compared with patterns of known immune cell types. Only the clusters that can be clearly annnotated were used for further experiments. Finally, we selected eight cell types (B cells, CD14 monocytes, NK cells, Treg, naive CD4 T cells, naive CD8 T cells, CD4 T effector memory and CD8 T effector memory) to perform deconvolution validation.

### Performance of DNN on different training datasets

We trained DNNs in DAISM-DNN on training datasets generated from expression data of purified cells to compare the performance of DNN with and without using real-life calibration samples. To this end, we generated two training datasets that used RNA-seq expression profiles of sorted cells and scRNA-seq data, and denoted them as “scRNAonly” and “RNAonly”, respectively. The generation of these training datasets followed the same linear mathematical operation as defined previously, with the only exception that real-life samples with ground truth cell fractions were not used in the mixing process. Briefly, for RNA-seq datasets, the expression of a simulated sample ***e*** was calculated as

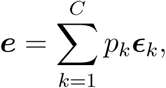

where *C* is the number of cell types involved in mixing, *p*_*k*_ (*k* = 1, …, *C*) are random variables with Dirichlet distribution that determined the fractions of different cells in the *in silico* mixed sample, and ***ϵ***_*k*_ is the expression profile of a randomly selected purified sample of cell type *k* from the respective RNA-seq or microarray dataset. For scRNA-seq dataset, ***e*** is given by

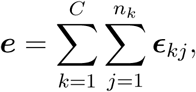

where *n*_*k*_ = 500 · *p*_*k*_ is the number of cells of type *k* extracted randomly from scRNA-seq datasets for mixing, and ***ϵ***_*kj*_ denote their expression profiles. Note that ***e*** were further TPM-normalized before used for training.

For comparison, we also generated two training datasets by DAISM with 200 randomly selected real-world samples from the SDY67 dataset as the calibration samples, which were further augmented using scRNA-seq or RNA-seq data of purified samples to train the DNN model in the DAISM-DNN pipeline. These two training sets were denoted as “DAISM-scRNA” and “DAISM-RNA”, respectively. The remaining 50 samples from SDY67 that were not used in creating the training set were used for testing. For t-SNE analysis, all the 50 test samples of SDY67 and 500 randomly selected artificial mixtures from each training datasets were plotted based on the common genes from SDY67 and the five training datasets. The parameter perplexity was set to 30 and the other parameters were set to their default values.

As suggested in Scaden, we also generated five training datasets that directly combined 5 different real-world bulk RNA-seq samples with simulated mixtures respectively. For fair comparison, the same number of training samples (16,000) were used in all training sets.

### Performance benchmarking

Since cell types’ abundances were resolved at different granularities in different deconvolution methods, regularizing the cell types of all methods to the same granularities has to be performed to facilitate a fair comparison. In this study, we only tested the performance of the benchmarked methods on six specific coarse-grain cell types (B cells, CD4 T cells, CD8 T cells, NK cells, monocytes, neutrophils) for comparison. The fine-grain cell type results of some methods were mapped to coarse-grain cell types according to the hierarchy of cell types defined in Strum et al.^21^.

CIBERSORT (CS) is a signature-based deconvolution algorithm which uses *ν*-SVR to estimate cell abundance. We obtained R code from the CIBERSORT website (https://cibersort.stanford.edu/). We used CIBERSORT with different signature matrices (LM22, IRIS, immunoStates and TIL10) and denoted them as four methods. The input data for CIBERSORT was in linear domain and all parameters were set to their default values. CIBERSORTx (CSx) is an extended version of CIBERSORT that generates a signature matrix from scRNA-seq data and provides two batch correction strategies (B-mode and S-mode) for cross-platform deconvolution. The B-mode was designed to remove technical differences between bulk profiles and signature matrices derived from bulk sorted reference profiles while S-mode was used for signature matrices derived from droplet-based or UMI-based scRNA-seq data. We experimented with both B-mode and S-mode. For B-mode, LM22 was used as the signature matrix. For S-mode, the scRNA-seq dataset PBMC8k was used to generate the signature matrix, which was further applied in deconvolution. We ran CIBERSORTx from its website (https://cibersortx.stanford.edu/). Quantile normalization was disabled when input was RNA-seq or scRNA-seq simulated mixture data. We used the R package *MuSiC* (https://github.com/xuranw/MuSiC) for MuSiC. MuSiC takes scRNA-seq data with cell type labels as reference. Deconvolution using MuSiC was performed with five coarse-grain cell types (B cells, CD4 T cells, CD8 T cells, NK cells and monocytes). The single cell PBMC dataset PBMC8k was used as reference data. ABIS enables absolute estimation of cell abundance from both bulk RNA-seq and microarray data. Deconvolution was performed through an R/Shiny app (https://github.com/giannimonaco/ABIS). The results output from ABIS were absolute cell frequencies and were divided by 100 in our study for comparison with other methods on RMSE. Scaden is a DNN-based deconvolution algorithm. We used the training datasets provided by Scaden (https://github.com/KevinMenden/scaden) which contains 32,000 artificial mixtures from four scRNA-seq PBMC datasets. Training was performed for 5000 steps per model on each dataset as recommended in the original paper. We ran quanTIseq, MCP-counter, EPIC, and xCell through R package *immunedeconv*^21^ which provided an integrated inference to benchmark on six deconvolution methods. The parameter *tumor* was set to FALSE when performing deconvolution on all PBMC datasets. As recommended in the original paper, EPIC was run with BRef as the signature set on PBMC samples.

### Statistical analysis

We used three evaluation criteria to compare the performance of DAISM-DNN methods. Pearson correlation *r* was used to measure the linear concordance between predicted cell proportions and the FACS ground truth. Lin’s CCC and the RMSE were further used to evaluate the performance for methods which enable absolute cell type proportion estimation.

Differences in continuous measurements were tested using the two-tailed Student’s t-test. Two-sided *p*-values were used unless otherwise specified, and a *p*-value less than 0.05 was considered significant. Ranking of the algorithms over multiple testing sets was determined using Friedman test with post hoc two-tailed Nemenyi test^35^. PRISM was used for basic statistical analysis and plotting (http://www.graphpad.com), and the Python or R language and programming environment were used for the remainder of the statistical analysis.

## Data availability

All expression datasets analyzed in this work, including accession codes and web links (if available), are listed in Supplementary Table S2.

## Code availability

The source code for DAISM-DNN is available at https://github.com/xmuyulab/DAISM-XMBD.

## Footnotes

1 XMBD: Xiamen Big Data, a biomedical open software initiative in the National Institute for Data Science in Health and Medicine, Xiamen University, China.

