## Supplementary Figures for "DAISM-DNN^XMBD^: Highly accurate cell type proportion estimation with *in silico* data augmentation and deep neural networks"

Y Lin et al.

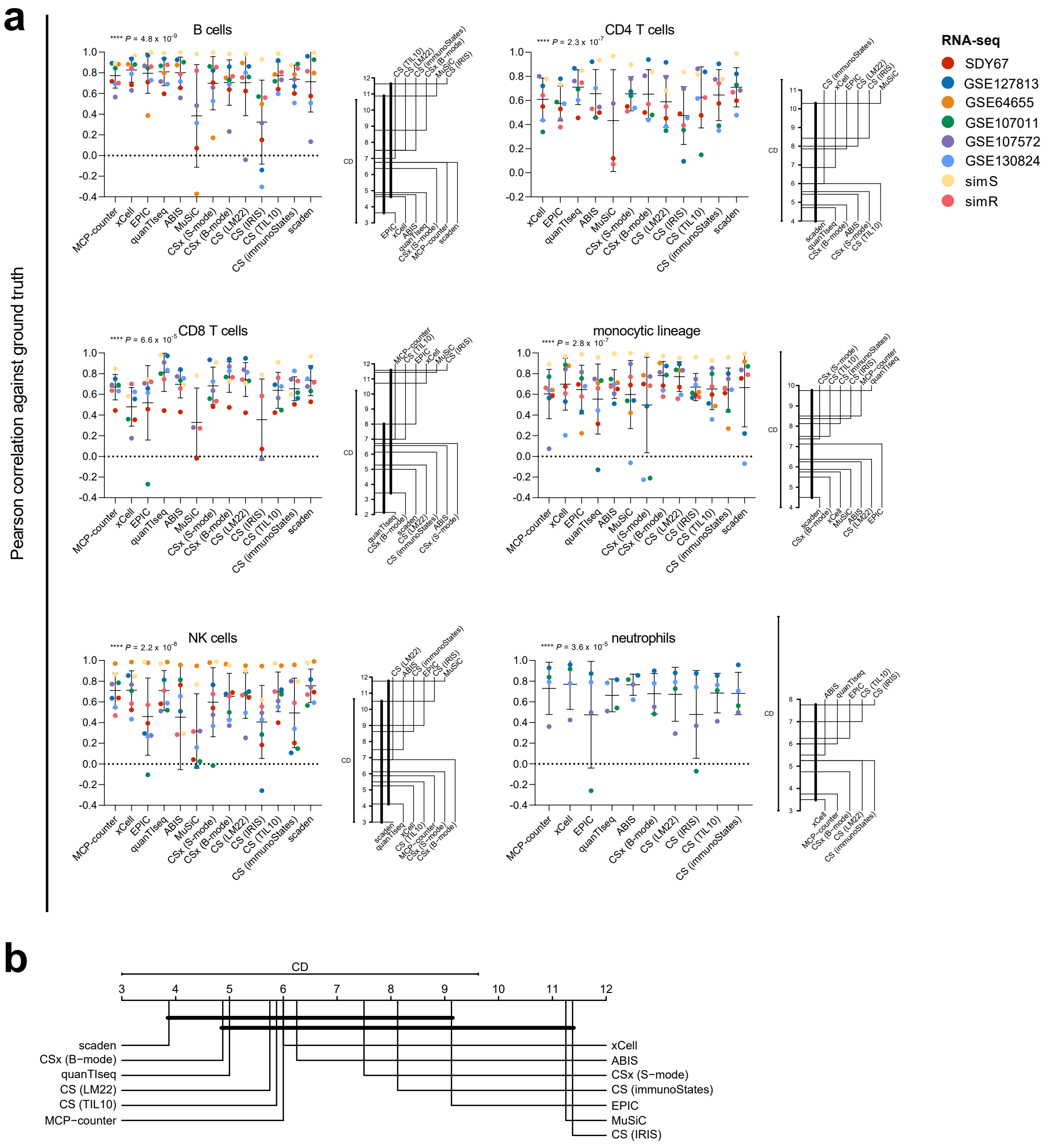


Figure S1: Evaluation of consistency in deconvolution performance across RNA-seq datasets. (a) Pearson correlations of the predicted cell type proportions of 13 deconvolution methods (including CIBERSORT using four different signature matrices) on six real-world datasets and two simulated datasets for six cell types, shown in scatter plots (left) with corresponding critical difference (CD) diagrams (right). Data are presented as means±SD. $*P < 0.05$, $**P < 0.01$, $***P < 0.001$, Friedman's rank sum test. (b) Comparison of 13 deconvolution methods on these eight datasets with post hoc two-tailed Nemenyi test. Groups of deconvolution methods that are not significantly different (at $P = 0.05$) are connected.

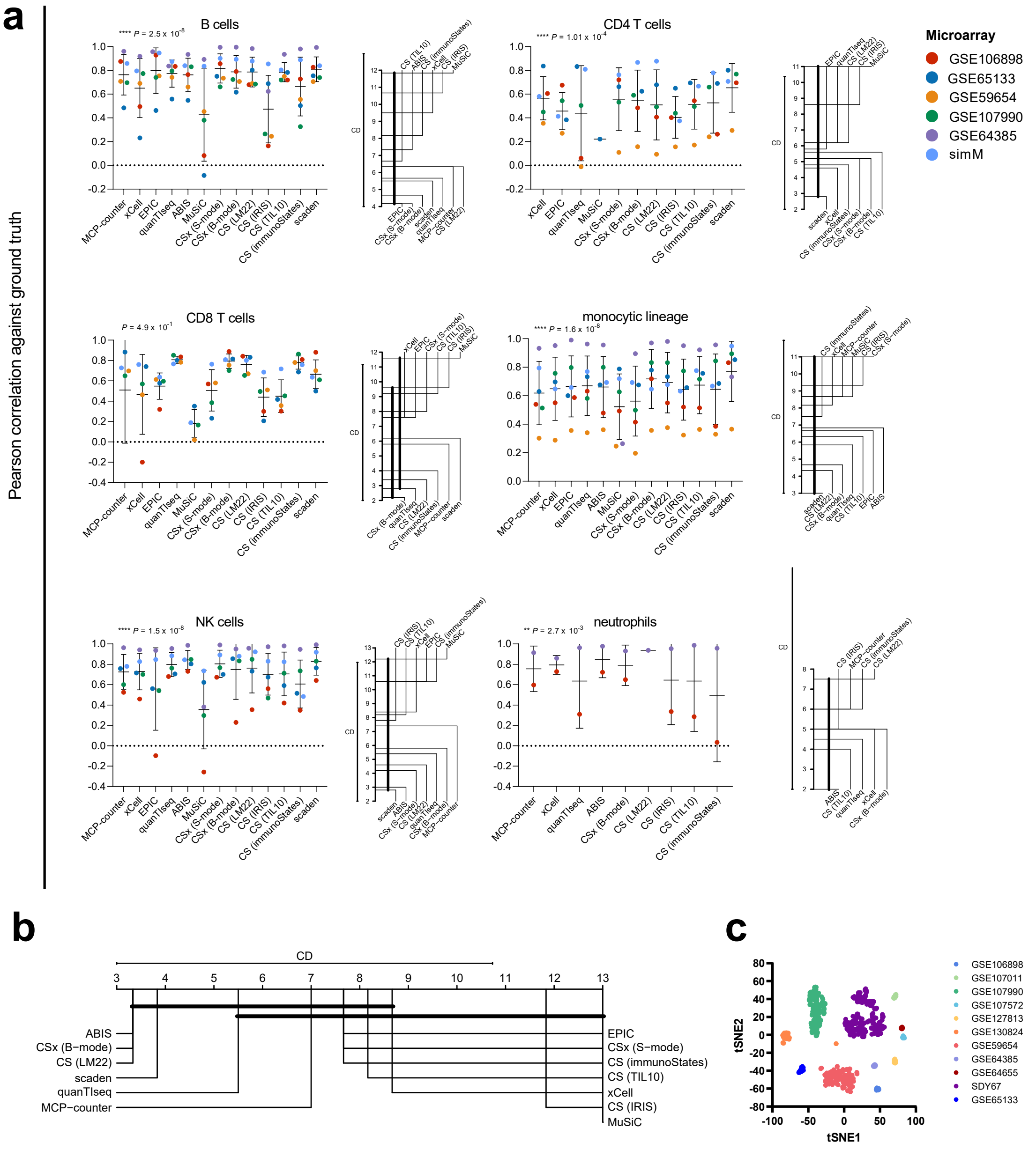


Figure S2: Evaluation of consistency in deconvolution performance across microarray datasets. (a) Pearson correlations of the predicted cell type proportions of 13 deconvolution methods (including CIBERSORT using four different signature matrices) on five real-world datasets and one simulated dataset for six cell types, shown in scatter plots (left) with corresponding critical difference (CD) diagrams (right). Data are presented as means±SD. *$P < 0.05$, **$P < 0.01$, ***$P < 0.001$, Friedman's rank sum test. (b) Comparison of 13 deconvolution methods on these six datasets with post hoc two-tailed Nemenyi test. Groups of deconvolution methods that are not significantly different (at $P = 0.05$) are connected. (c) t-SNE projection of eleven real-world datasets. Each dot represents one single sample, coloured by datasets.

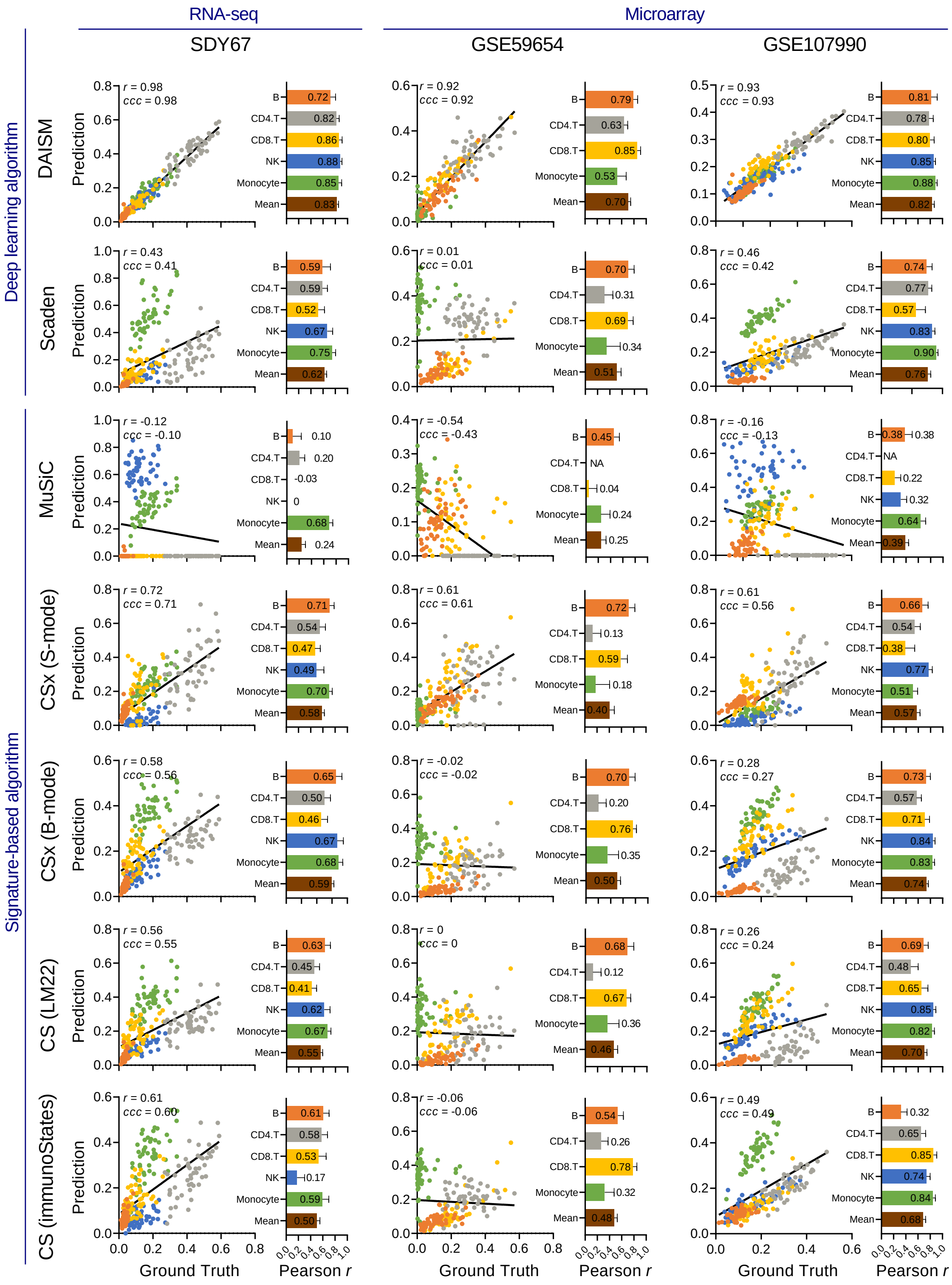
RNA-seq
Microarray
SDY67
GSE59654
GSE107990
Deep learning algorithm
Signature-based algorithm
DAISM
Scaden
MuSiC
CSx (S-mode)
CSx (B-mode)
CS (LM22)
CS (immunoStates)
Prediction
Ground Truth
Pearson r
B
CD4.T
CD8.T
NK
Monocyte
Mean
r = 0.98
ccc = 0.98
0.72 0.82 0.86 0.88 0.85 0.83
r = 0.92
ccc = 0.92
0.79 0.63 0.85 0.53 0.70
r = 0.93
ccc = 0.93
0.81 0.78 0.80 0.85 0.88 0.82
r = 0.43
ccc = 0.41
0.59 0.59 0.52 0.67 0.75 0.62
r = 0.01
ccc = 0.01
0.70 0.31 0.69 0.34 0.51
r = 0.46
ccc = 0.42
0.74 0.77 0.57 0.83 0.90 0.76
r = -0.12
ccc = -0.10
0.10 0.20 -0.03 0 0.68 0.24
r = -0.54
ccc = -0.43
0.45 NA 0.04 0.24 0.25
r = -0.16
ccc = -0.13
0.38 NA 0.22 0.32 0.64 0.39
r = 0.72
ccc = 0.71
0.71 0.54 0.47 0.49 0.70 0.58
r = 0.61
ccc = 0.61
0.72 0.13 0.59 0.18 0.40
r = 0.61
ccc = 0.56
0.66 0.54 0.38 0.77 0.51 0.57
r = 0.58
ccc = 0.56
0.65 0.50 0.46 0.67 0.68 0.59
r = -0.02
ccc = -0.02
0.70 0.20 0.76 0.35 0.50
r = 0.28
ccc = 0.27
0.73 0.57 0.71 0.84 0.83 0.74
r = 0.56
ccc = 0.55
0.63 0.45 0.41 0.62 0.67 0.55
r = 0
ccc = 0
0.68 0.12 0.67 0.36 0.46
r = 0.26
ccc = 0.24
0.69 0.48 0.65 0.85 0.82 0.70
r = 0.61
ccc = 0.60
0.61 0.58 0.53 0.17 0.59 0.50
r = -0.06
ccc = -0.06
0.54 0.26 0.78 0.32 0.48
r = 0.49
ccc = 0.49
0.32 0.65 0.85 0.74 0.84 0.68

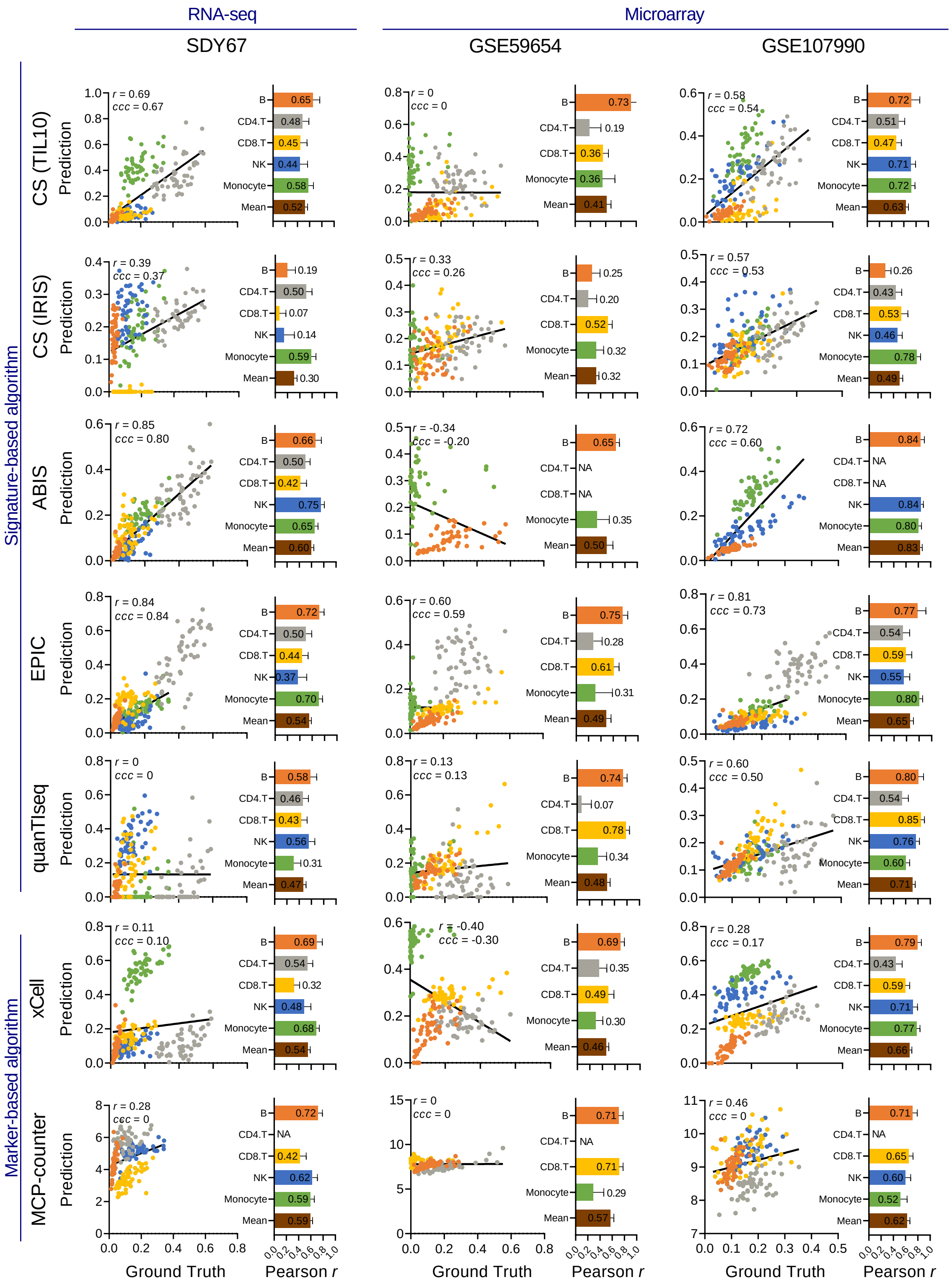


Figure S3: Performance evaluation of DAISM-DNN and other 12 deconvolution methods, tested on one RNA-seq dataset SDY67 (left) and two microarray datasets GSE59654 (middle) and GSE107990 (right). In each subplot, the scatter plot shows a global Pearson correlation ($r$) and concordance correlation coefficient (CCC). The bar plot displays per-cell-type Pearson correlation. The value in bar plots indicates the mean value over 30 experiments. Since different methods provide fraction of different cell types, we only validated the ones detected from flow cytometry in each dataset. NAs in the bar plots indicate cell types that cannot be predicted by this method.

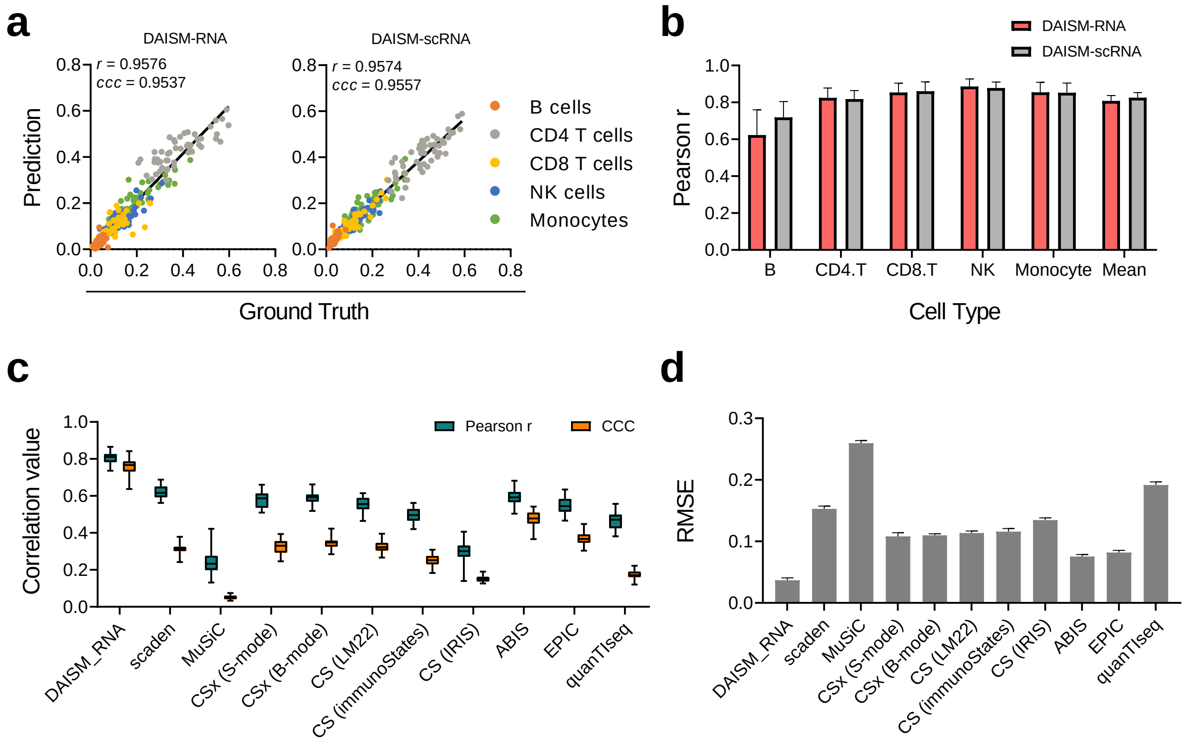


Figure S4: Performance evaluation of DAISM-RNA mode on RNA-seq dataset SDY67. (a-b) Deconvolution performance comparison between DAISM-RNA and DAISM-scRNA mode. Global Pearson correlation ($r$) and CCC are shown in scatter plots (a), and per-cell-type Pearson correlation are shown in the bar plot (b). (c-d) Comparison of DAISM-RNA mode with other methods by mean of per-cell type Pearson correlation and CCC (c) and RMSE (d). All data in bar plots are presented as the mean±SD.

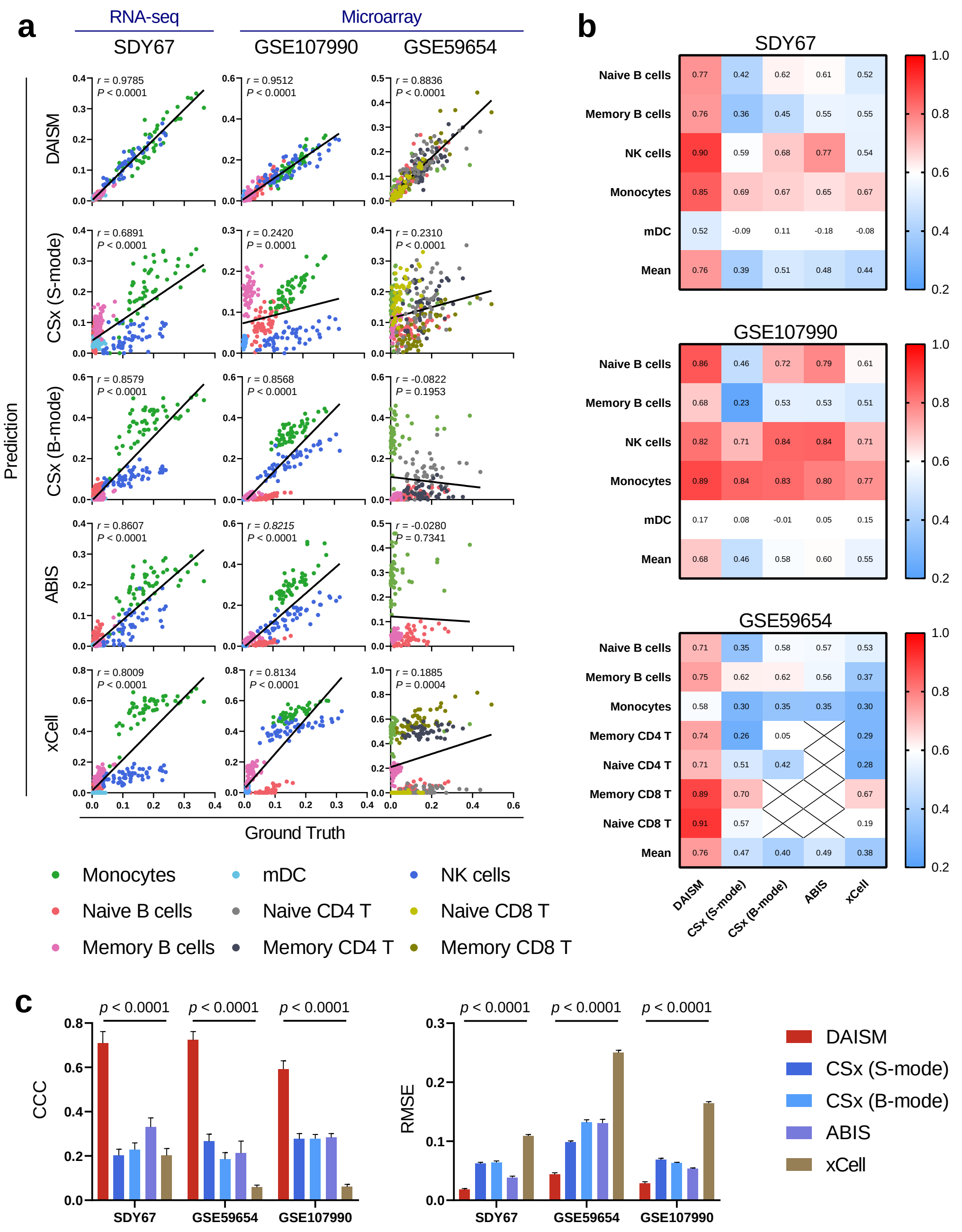


Figure S5: Performance of DAISM-DNN in fine-grain mode and benchmarking with four methods, validated using one RNA-seq dataset SDY67 and two microarray datasets GSE107990 and GSE59654. (a) Global Pearson correlation ($r$) between predicted and ground truth cell fraction. SDY67 and GSE107990 have five cell types: naive/memory B cells, NK cells, monocytes, myeloid dendritic cells (mDC), and GSE59654 has seven cell types: naive/memory B cells, monocytes, naive/memory CD4/CD8 T cells. (b) Heatmap summarizing performance of all methods after 30 bootstrapping experiments by mean of Pearson correlation on each cell type and the mean of per-cell-type Pearson correlation (the last row in each heatmap). (c) Barplots of RMSE (right) and CCC (left) for five methods. Data are presented as means±SD. Two-sided paired Student's t tests were used to compare DAISM-DNN with other methods.

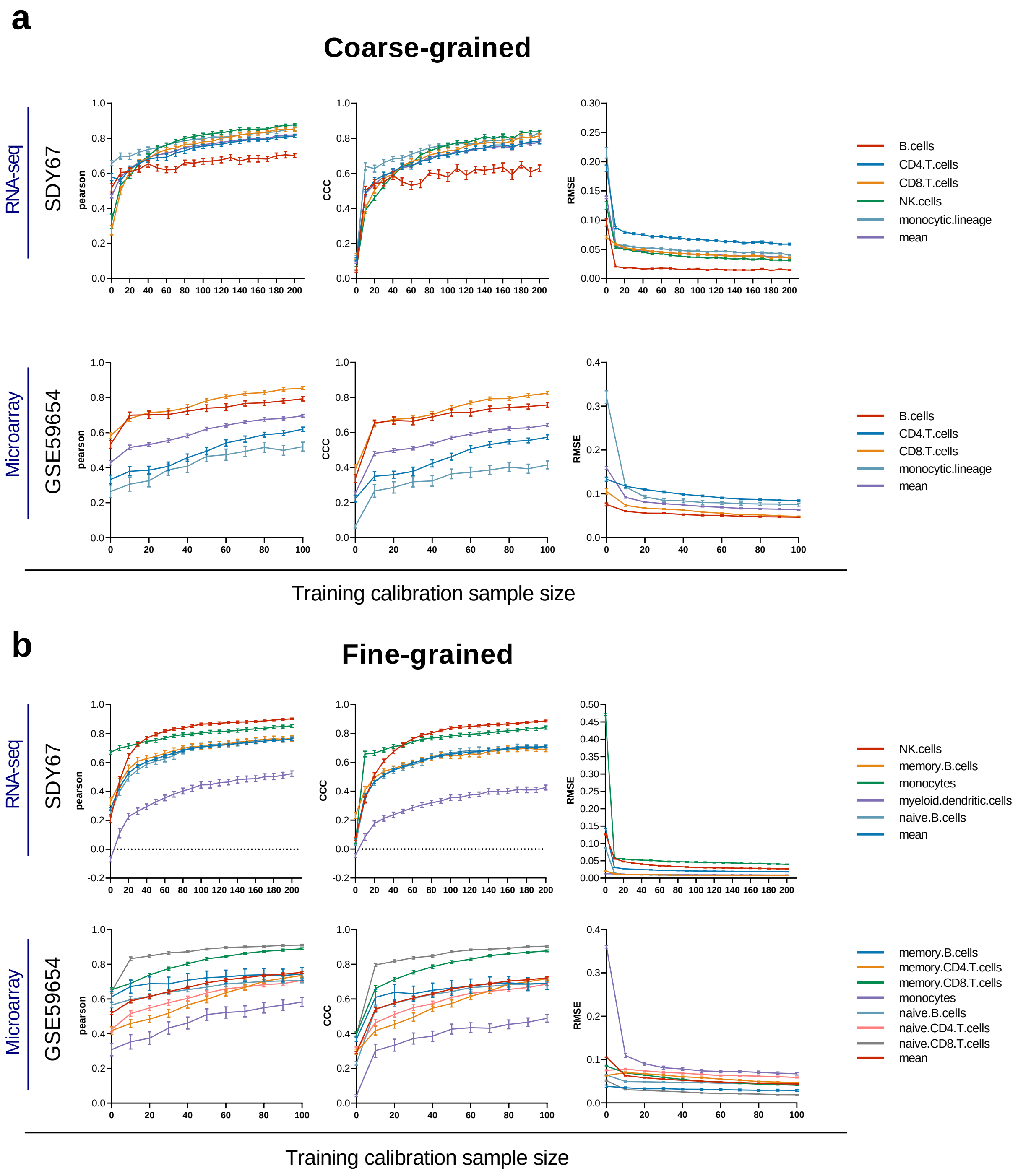


Figure S6: The effect of calibration sample size on the DAISM-DNN pipeline, assessed by the Pearson correlation, CCC and RMSE of the cell type proportion estimation results. (a) We used four scRNA-seq datasets as augmentation data to generate simulated training data, with different calibration sample size of RNA-seq dataset SDY67 and microarray dataset GSE59654 for coarse-grain cell types. Data are presented as means±SEM, coloured by different cell types in each dataset. (b) Results for fine-grain cell types.

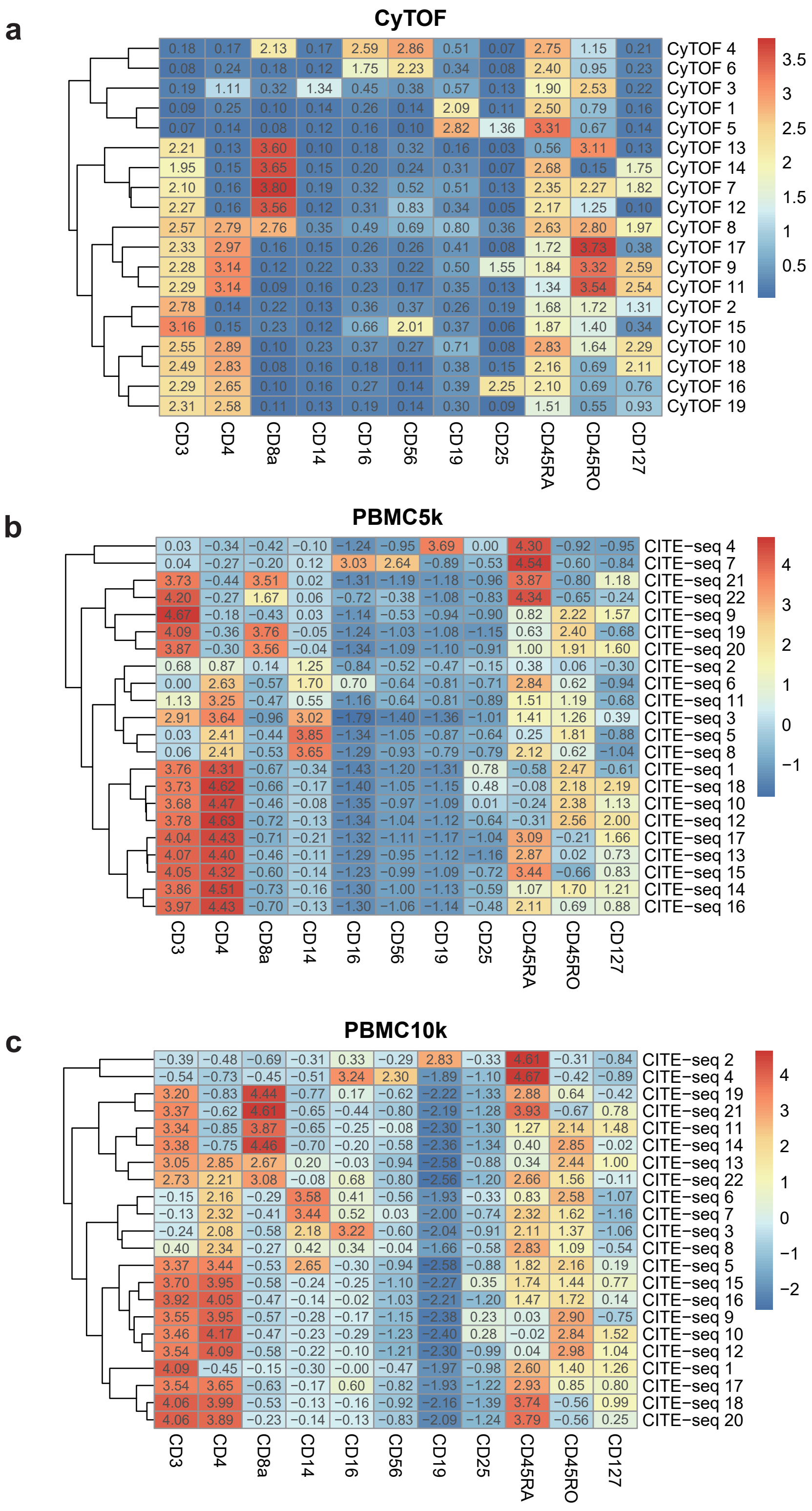


Figure S7: Heatmap showing mean values of normalized markers expression in each Phenograph clusters. CyTOF in-house dataset (a) and two public CITE-seq datasets (b) PBMC5k and (c) PBMC10k.

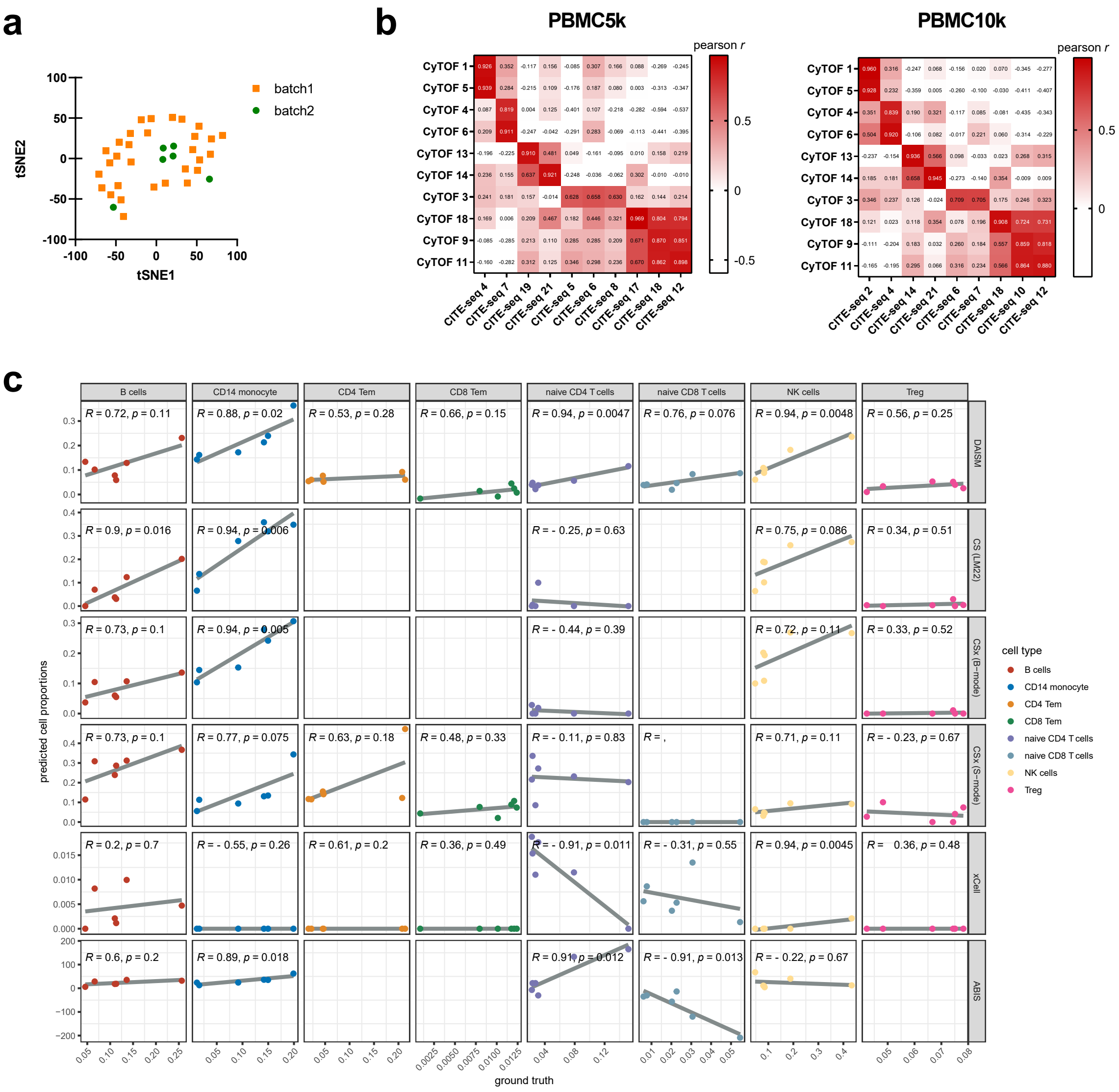

Figure S8: DAISM-DNN enables cross-batch accurate cell proportion estimation. (a) The t-SNE projection of in-house dataset colored in different batches. (b) Heatmap showing pearson correlation between CyTOF data clusters and CITE-seq data clusters used for augmentation. (c) Scatterplots for eight immune cell types of ground truth (x axis) and predicted values (y axis) for DAISM-DNN, CSx (including two batch-correction modes), xCell and ABIS on in-house data. Numbers inside the plotting area signify Pearson correction values and p-values (Student's t-test).
